# Advancing Mobile Neuroscience: A Novel Wearable Backpack for Multi-Sensor Research in Urban Environments

**DOI:** 10.1101/2025.04.13.648607

**Authors:** João Amaro, Rafael Ramusga, Ana Bonifácio, João Frazão, André Almeida, Gonçalo Lopes, Ata Chokhachian, Daniele Santucci, Paulo Morgado, Bruno Miranda

## Abstract

The rapid global urbanisation has intensified the need to understand the complex interactions and impacts that city environments have on human physical or mental health and well-being. Traditional indoor laboratory-based approaches conduct experiments in well controlled settings but, while advantageous for their controlled conditions, they often lack ecological validity. To address this gap, we present the “eMOTIONAL Cities Walker Backpack” — a wearable unit developed for synchronously collecting multi-modal data in dynamic real-world settings. Designed for both indoor and outdoor use, the backpack integrates environmental (for microclimate, air pollution and noise) and physiological sensors (including electroencephalography, eye-tracking and wrist-based biosensors for cardiovascular monitoring) to enable the study of human experience in naturalistic urban environments. In this paper, we describe the technical specifications and implementation of this technology during outdoor acquisitions across selected urban locations in the city of Lisbon. We also highlight its potential for methodological comparison with traditional lab-based tasks (particularly through the use of equivalent sensing technologies), and thus advancing the field of translational research in mental health and urban studies.

## 1. Introduction

The 21st century has been identified as the *Century of the City*, highlighting a significant global shift toward urban living [1]. According to the United Nations (UN-DESA, 2024), the global population is expected to continue to increase for another five to six decades, reaching a peak of approximately 10.3 billion people by 2080 [2]. Urban development has yielded positive outcomes by generating new opportunities and enhancing overall well-being and quality of life; nevertheless, it can also lead to adverse consequences, negatively impacting people’s health and well-being [3].

While it is well-documented that urban residents are more susceptible to experiencing symptoms of anxiety and depression compared to their rural counterparts [4], emerging evidence suggests that exposure to certain environments – including highly vegetated spaces or those with bodies of water, may provide psychological restoration in both metropolitan and countryside settings [5–8]. However, the biological mechanisms underlying these observations remain largely unclear, with only a limited number of studies investigating the brain and body physiological responses that causally link specific environmental exposures to physical or mental health outcomes [9]. Moreover, such research on human-environment interactions have predominantly involved stationary participants in controlled laboratory settings; although recent advances in neuroscience technology have enabled the recording of human brain activity during freely moving behaviours [10].

The emerging field of *Neurourbanism*, also referred to as *Environmental Neuroscience*, embraces an interdisciplinary framework that integrates neuroscientific techniques with traditional assessment methods to achieve a more objective and comprehensive understanding of human responses to both built and natural urban environments [11,12]. Central to this approach is the investigation of how specific urban features influence subjective experience and psychological well-being through direct measurement of brain activity. Recent years have seen a growing body of research employing mobile neuroimaging tools – particularly mobile electroencephalography (EEG) – to explore these dynamics *in situ* [13]. By combining various sensor-based technologies, this holistic methodology offers deeper insights into the complex interplay between environmental context and mental health, with the potential to inform evidence-based urban planning and public health strategies [14–16].

Overall, contemporary research work should prioritize enhancing ecological validity by examining human physiological and psychological responses as they naturally occur during daily interactions within the city [17]. Adopting this approach enables researchers to capture the complexity and authenticity of physiological and cognitive processes in realistic contexts, thereby enhancing the generalizability and applicability of findings. This improved ecological perspective offers significant practical value, providing policymakers, urban planners, and healthcare professionals with robust insights that directly inform effective urban design and public health interventions. Nonetheless, this implies methodological challenges –such as the adaptation of multisensory and dynamic naturalistic paradigms (e.g., a city walk experience) to generate meaningful and valid results (that could be related to the ones obtained in well-controlled laboratory settings; see [18,19]); and technical hurdles – combine sensors that synchronously capture and track mobile body and brain responses, as well as the respective environmental exposures (e.g., the visual and noise stimuli, air quality or microclimate).

### 1.1. Wearable sensors and outdoor studies

Simulations based approaches to the influence of comfort on cities [20] usually underestimate the human dimension. Human behaviour is dependent on environmental context, and it is known that human actions in an urban environment tend to optimize human comfort [21]. Using mobile sensors to assess environmental conditions [21–24] allows the understanding of the interplay between climate and its effect on human behaviour [25].

There is a wealth of wearable sensors created for environmental monitoring, such as those for thermal, visual, acoustic, and air quality environmental effects. While most studies consider wearable sensors that span these environmental factors, seldom were these integrated with biometric data [26]. On the other end, the majority of research relying on health monitoring for environmental exposure still relies heavily on subjective measurements (such as self-report questionnaires), and only few utilize portable health sensors [27]. The vast majority of studies monitoring the human body focus on cardiovascular measures and lung function as a function of air quality [e.g., exposure to particulate matter, or PM)], without much focus on other relevant physical environmental factors such as noise and temperature [26,27]. Notwithstanding, efforts have been made to facilitate the simultaneous use of environmental sensors with biosensing — for example, one study gathered meteorological data through a wearable sensor station, as well as simultaneously skin temperature monitoring to assess thermal comfort [21].

Crucially, the assessment of mental states in urban context and its effect on mental health necessitates the use of brain imaging methods [28]. For assessing brain activity during outdoor walks, it is either used the EEG or a functional near-infrared spectroscopy (fNIRS) technologies [13]. As EEG is capable of millisecond temporal resolution alongside a better portability, it is often preferred over fNIRS [29]; the EEG has also become widely available (specially with the emergence of consumer-grade EEG) for non-laboratory settings and for application in brain-computer interfaces (BCI) [19,30]. Such increase in available devices has also fostered a wealth of studies focused on the performance of EEG (and its signal quality) in moving subjects; as well as reflections about its correct use in these settings [31–33]. Attempts to use mobile EEG studies for more naturalistic evaluation of the surrounding environment, combine laboratory-based settings with, for example, a treadmill [34]; or ask participants to go for a walk with a wireless EEG device (connected to a computer or smartphone often carried by the participant) — but very often using EEG systems with fewer channels or less robust.

The drive for ecological validity in experimental designs has been driving interesting advances in human cognitive neuroscience, at the same time as it ameliorates the recording techniques and methodologies to conduct experiments in outdoor settings [18]. One challenge to this potential paradigm shift is the need to control for a vast number of influencing variables — as in the case for complex and dynamic urban environments. This gap can be addressed by creating a wearable data collection unit capable of synchronous acquisition of a multitude of data types — i.e., both human responses and environmental exposures.

### 1.2. Scope and Objectives

The present work was conducted as part of the eMOTIONAL Cities project (https://emotionalcities-h2020.eu/) – an interdisciplinary European Union Horizon 2020 Research and Innovation Action aiming to elucidate how natural and built urban environments influence human cognitive and emotional processes, particularly through their neurobiological underpinnings. The project has integrated advanced neuroscience methodologies, including neuroimaging and physiological monitoring, to gather robust evidence about the interactions between urban characteristics and individual well-being.

Here, we aim to report the methodological foundations for neuroscience research on free-moving humans in urban environments. It is introduced a new wearable experimental platform that combines multiple environmental and health sensors, suitable for both indoor and outdoor neuroscience experiments in natural urban settings. We detail the implementation of a flexible experimental framework that ensures comparability and reliability of data across indoor and outdoor paradigms.

By bridging lab-based and field-based approaches, this work contributes to the growing body of evidence on how urban spaces influence cognitive and emotional processes. It also highlights the necessity of validated, ecologically relevant methods for capturing the complexity of human experiences in the city, ultimately supporting evidence-based urban planning aimed at promoting well-being.

As such, this paper focuses on the description of a novel multimodal wearable apparatus combining sensors for environmental factors with biosensing (including the use of EEG for better mental state assessment characterisation) that could be used in naturalistic settings.

## 2. Materials and methodology

### 2.1. Participants

Participants were recruited using convenience sampling through serendipitous methods (to identify and invite eligible individuals), including outreach to personal and professional networks, such as colleagues, friends, and acquaintances. They were required to be 18 years of age or older, residing in Lisbon, fluent in Portuguese or English and have no history of major psychiatric or neurological disorder. Participants were invited to participate in both indoor and outdoor experiments; but only 3 were involved in both. Each participant was free to repeat the experiment on other paths, up to a maximum of three completed paths per participant. As such, 43 participants contributed to a total of 60 sessions (10 sessions per path). Participants received a voucher as compensation for their time and involvement. All participants provided informed consent, and the study was approved by the Ethics Committee of the Academic Centre of Medicine of Lisbon.

### 2.2. Experimental procedures and stimuli

Participants were randomly assigned to one of six possible paths, which were chosen based on urban areas of interest in the city of Lisbon, Portugal. These urban areas were defined in a workshop which reunited stakeholders and professionals to determine a set of city paths with a wide variety of urban characteristics in the city of Lisbon for data collection [35]. Prior to beginning the task, participants were fitted with the multi-modal data acquisition backpack apparatus (described below); and instructions to the participants emphasized the need to abstract away from the experimental context in order to simulate a common daily walk as done in normal contexts. The task consisted of a walk on a predefined path (around 1 kilometre long), with the researchers following behind the participant and giving verbal instructions when necessary. Each path had a specific amount of “checkpoints”, which represented areas that were filmed and shown in a lab-based EEG task, with the purpose of comparing and validating data from both paradigms (full analysis of this comparison will be considered in a separate work). Upon reaching the checkpoint, three sequential actions were asked from the participant: observing the environment while standing still; walking for 20 seconds; and finally, answering questions to assess the environment’s level of naturalness, crowdedness, as well as elicited feelings of valence and arousal (**Figure 1**).

**Figure 1.**
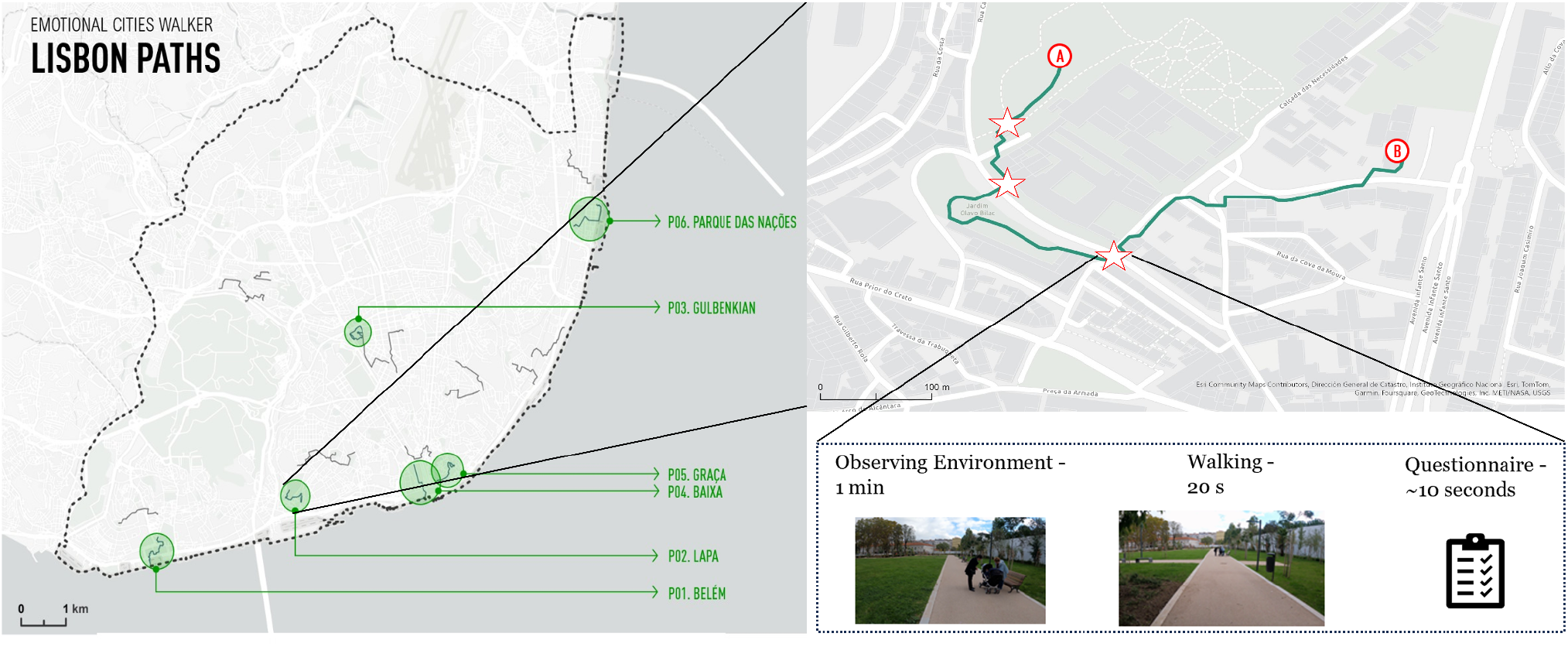
The outdoor walking task was conducted across six different paths in the city of Lisbon, which were chosen based on certain geographical and sociodemographic characteristics. Between the start (A) and end (B), each path also had some “checkpoints” (marked with stars) – where participants were asked to pause (observing the surroundings still for one minute), walk for 20 seconds and then pause again to answer several questions about the walking experience. The data gathered at these checkpoints was then compared to data acquired in a related laboratory-based experiment that used first-person video walks of the same section of the path.

## 3. Results

### 3.1. eMOTIONAL Cities Walker backpack

The wearable data collection unit, named *eMOTIONAL Cities Walker*, is an ergonomically-driven apparatus in the shape of a backpack whose function is to acquire several different streams of data from its user and the surrounding environment (**Figure 2**). This was achieved using a portable, mobility-focused computer—the HP® VR Backpack PC(HP Inc., Palo Alto, CA, USA)—which is designed to fit into a backpack-style frame, allowing participants to comfortably wear it during outdoor walks. The dual external batteries ensured continuous operation in the field for up to two hours. Its compact design and wireless connectivity made it suitable for unobtrusive outdoor use with multiple sensors. To facilitate the interaction between the researcher and the computer a touchscreen interface was added, allowing for a smooth implementation and supervision of the scientific task. **Table 1** provides a summary of the sensors included in the equipment, listing the sensor name, brand, measured parameters, units, and data collection frequency.

**Table 1.**
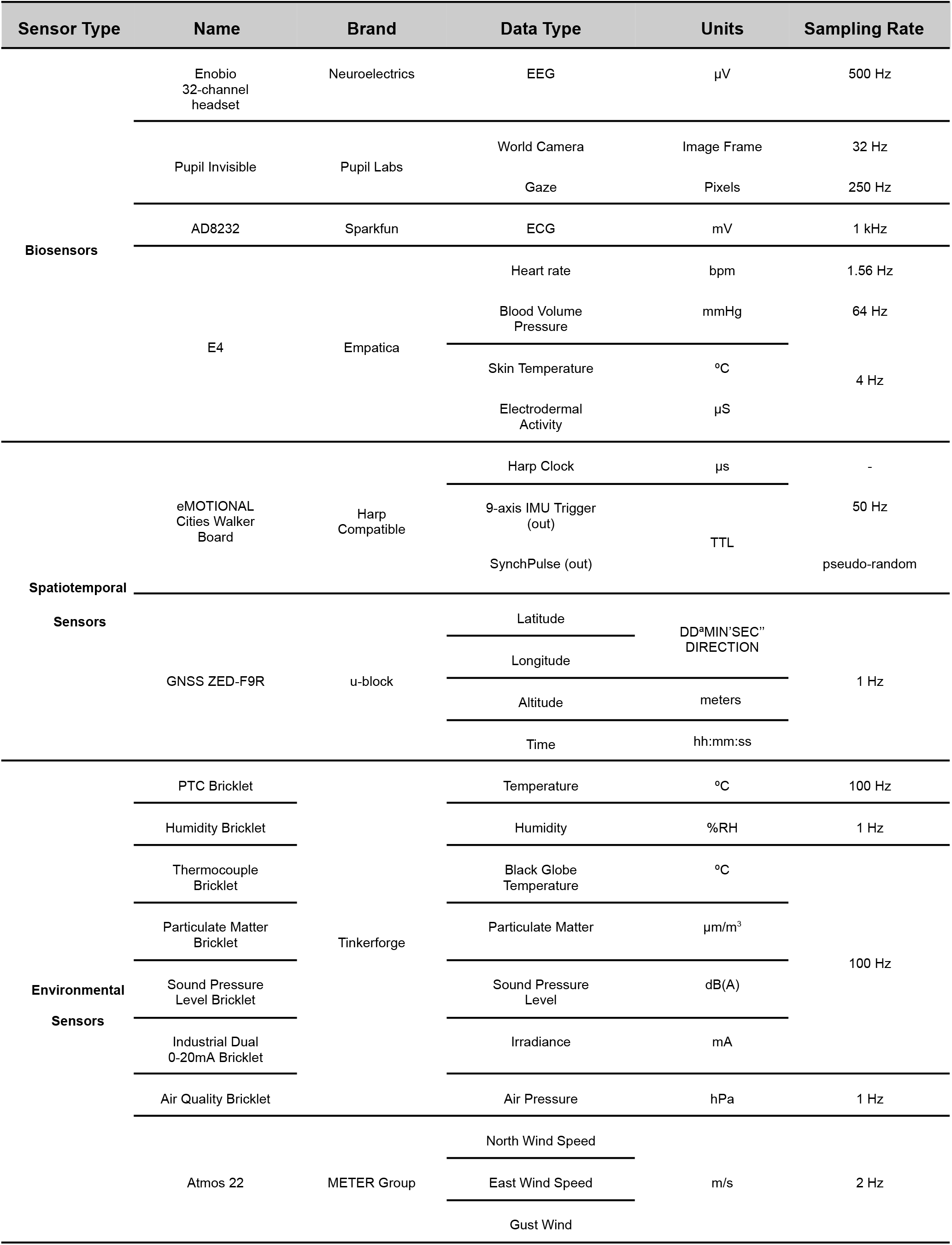
Summary of the sensors present in the eMOTIONAL Cities Walker.

**Figure 2.**
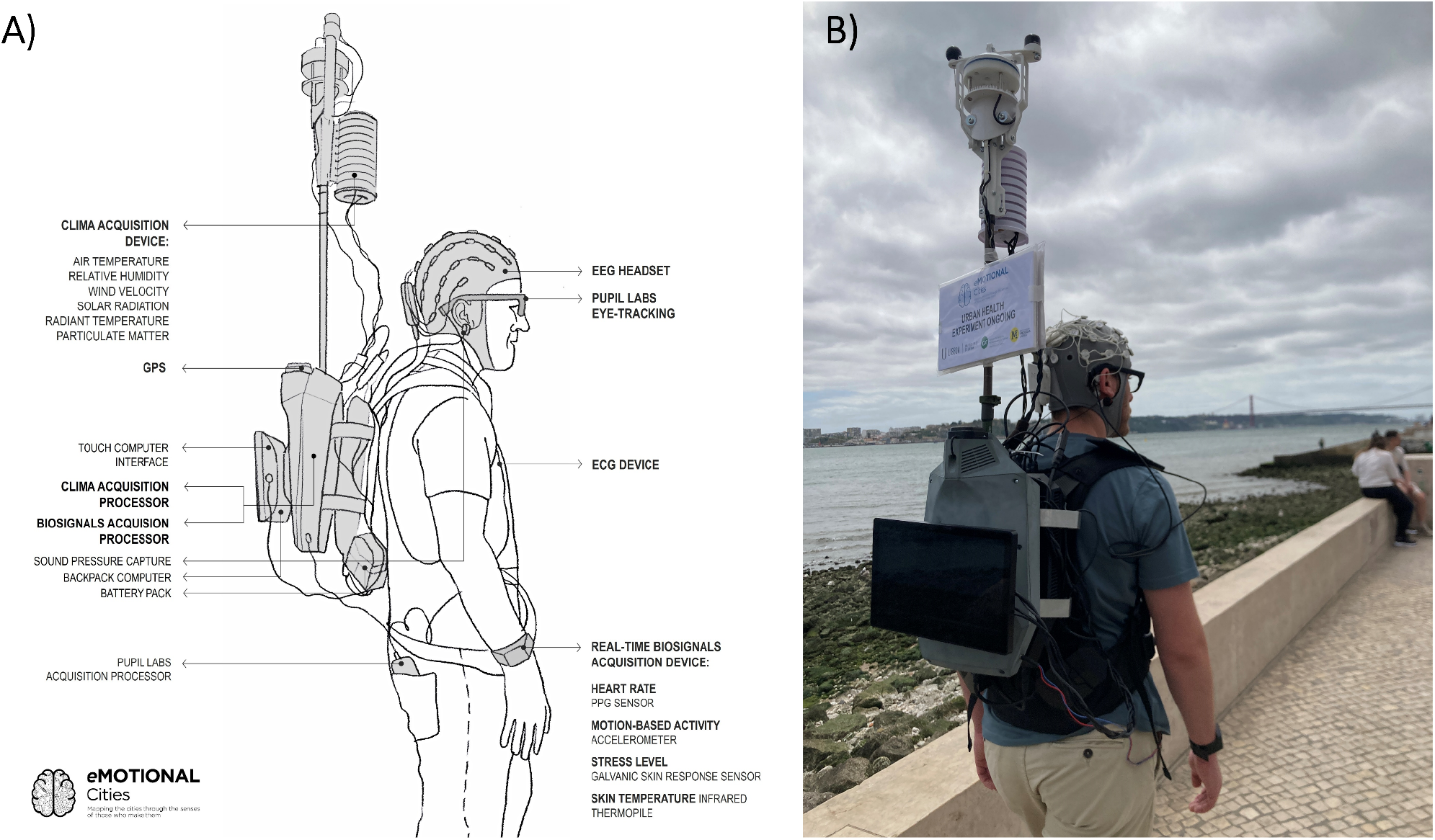
**A)** Schematic of the eMOTIONAL Cities Walker Backpack wearable data collection unit with indication of the devices and sensors. **B)** Side view photo of a participant wearing the eMOTIONAL Cities Walker Backpack.

### 3.2. Sensors for human physiology and behavioural signals

#### Electroencephalogram (EEG)

EEG data is acquired using the Enobio® (Neuroelectrics, Barcelona, Spain) system, in combination with a neoprene headcap, electrode cable sets for 32 channels, and dry EEG electrode technology (Drytrodes)– which allows the collection of data quickly and comfortably, with no traces of gel on the participant’s head [19]. Event information was designed to be sent via lab streaming layer (LSL) (https://github.com/sccn/labstreaminglayer) to allow synchronisation with the EEG data.

#### Peripheral physiological signals

Physiological data were collected using the E4 wristband (Empatica Inc., Boston, MA, USA) – a wristband that allows for data derived from: Photoplethysmogram (PPG) – allowing for blood volume pulse (BVP), inter beat interval (IBI) and heart rate (HR); electrodermal activity (EDA) expressed in microsiemens; 3-axis accelerometer (acceleration ranging from −2g to 2g); and skin temperature (expressed on the Celsius scale) sensors.

#### Electrocardiogram (ECG)

For the collection of the ECG data, the sparkfun AD8232 board was used. This sensor provides more reliable data than the in-built ECG from Empatica’s wristband, which requires minimal movement of the wrist for a good quality signal, a requirement impossible to meet for an outdoor walk-based task. Furthermore, the usage of specific sticker ECG sensors, in this case, Ambu Blue Sensor, provides a gel interface between the skin and the ECG electrode, which translates into a cleaner signal.

#### 9-axis Inertial Measurement Unit (IMU)

For the collection of movement data, it is used the BNO055 Intelligent 9-axis absolute orientation sensor. We can measure (with more accuracy and detail than GNSS) the sudden subject accelerations, changes in direction, resting or stopping periods.

#### Eye-tracking (ET)

Eye-tracking data were collected using Pupil Invisible glasses (Pupil Labs GmbH, Berlin, Germany) – a device that allows the combination of eye tracking data and an external camera, leading to a geotagged “egocentric” perspective of a subject walking in the city. The analysis of gaze could isolate elements in the external environment capturing the attention of the user at any point in time; and could be combined with object recognition techniques to group landmarks or areas/objects of interest (in both space and time).

### 3.3. Environmental sensors

The collection of environmental data was conducted using sensors from Tinkerforge and METER Group. The Tinkerforge sensors measured various environmental parameters, including humidity with the Humidity Bricklet, black globe temperature with the Thermocouple Bricklet, particulate matter (μg/m^3^) with the Particulate Matter Bricklet, sound pressure level (dbA) with the Sound Pressure Level Bricklet, irradiance (W/m^2^) with the Industrial Dual 0-20 mA Bricklet, and air pressure (hPa) with the Air Quality Bricklet. In addition, air temperature, wind velocity, and direction were measured using the ATMOS 22 sensor from METER Group.

### 3.4. Spatiotemporal Sensor Synchronization

Temporal synchrony between data streams is achieved with the eMOTIONAL Cities Walker’s motherboard, a device compatible with HARP, which is a standard for asynchronous real-time data acquisition and experimental control in neuroscience. Spatial data is acquired with the Global Navigation Satellite System (GNSS) using the ZED-F9R high precision dead reckoning module from u-blox. For each GNSS position data point, there is an associated high precision Coordinated Universal Time (UTC) timestamp. This timestamp is later used to give spatiotemporal context to all data from the other sensors.

### 3.5. Integration software

For synchronization of different data types, both temporally and spatially, all data collection has been integrated into a common acquisition framework, based on the open-source Bonsai programming language, for data streams, using standard interfaces to access each data source [36]. For an overview depicting the different data sensors and communication protocols with the bonsai software, see **figure 3**.

**Figure 3.**
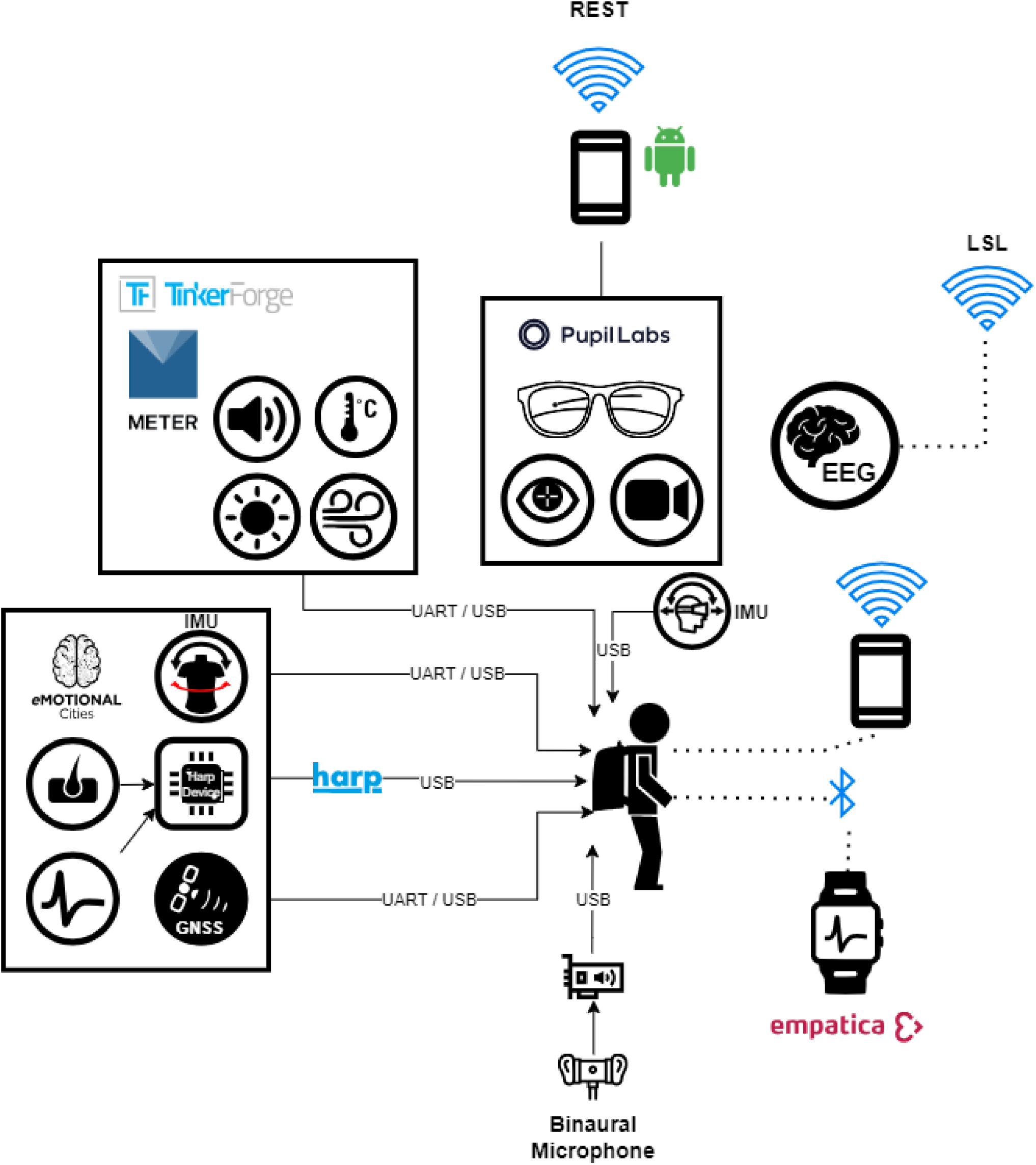
Integration of the different data streams at the software level.

Bonsai generates random timing events that the eMOTIONAL Cities Walker’s board adds as a Harp timestamp. These Harp timestamps are then routed as a time-to-live (TTL) to synchronize with other sensors capable of hardware-level synchronization, such as TinkerForge, Enobio’s EEG system, and the u-blox GNSS. The 9-Axis IMU sampling is triggered and Harp-timestamped by the eMOTIONAL Cities Walker board. For sensors lacking hardware synchronization, a software approach is pursued by fetching the most recent Harp timestamp and using it used to tag Empatica’s and Pupil Labs’ data as they arrive. The same approach is also used to tag the beginning of the experiment and other relevant events of the experimental protocol, which is done manually by the researcher. The random timing event and the harp timestamp are used to temporal align every sensor data stream and even recover synchronization if data or events are lost. Data is thus aligned with GNSS position and UTC high precision timestamps.

### 3.6. Sensor Reliability

Overall, the eMOTIONAL Cities Walker Backpack proved successful in its function as a reliable apparatus for outdoor experiments. The setup time for equipping each participant was approximately 10 minutes, which is relatively efficient considering the complexity of the equipment involved. Participants across different age groups successfully completed the 25-minute walking protocol with minimal difficulty, indicating that the ergonomic design of the backpack was largely effective. Despite the overall success in data collection, the fidelity of sensor recordings varied across the 60 acquisition sessions in Lisbon. **Figure 4** illustrates the recording status of each sensor during these sessions. The integration of multiple sensors, mostly connected via cables, was dependent upon the integrity of the connection, whose probability of failure increases due to wear and tear over repeated use. Additionally, connections are susceptible to spurious cable disconnections and signal interference, especially in mobile settings where movement can cause strain on connectors and cables. These reasons underlie the selective failure of some sensors to capture data completely or only partially during certain sessions.

**Figure 4.**
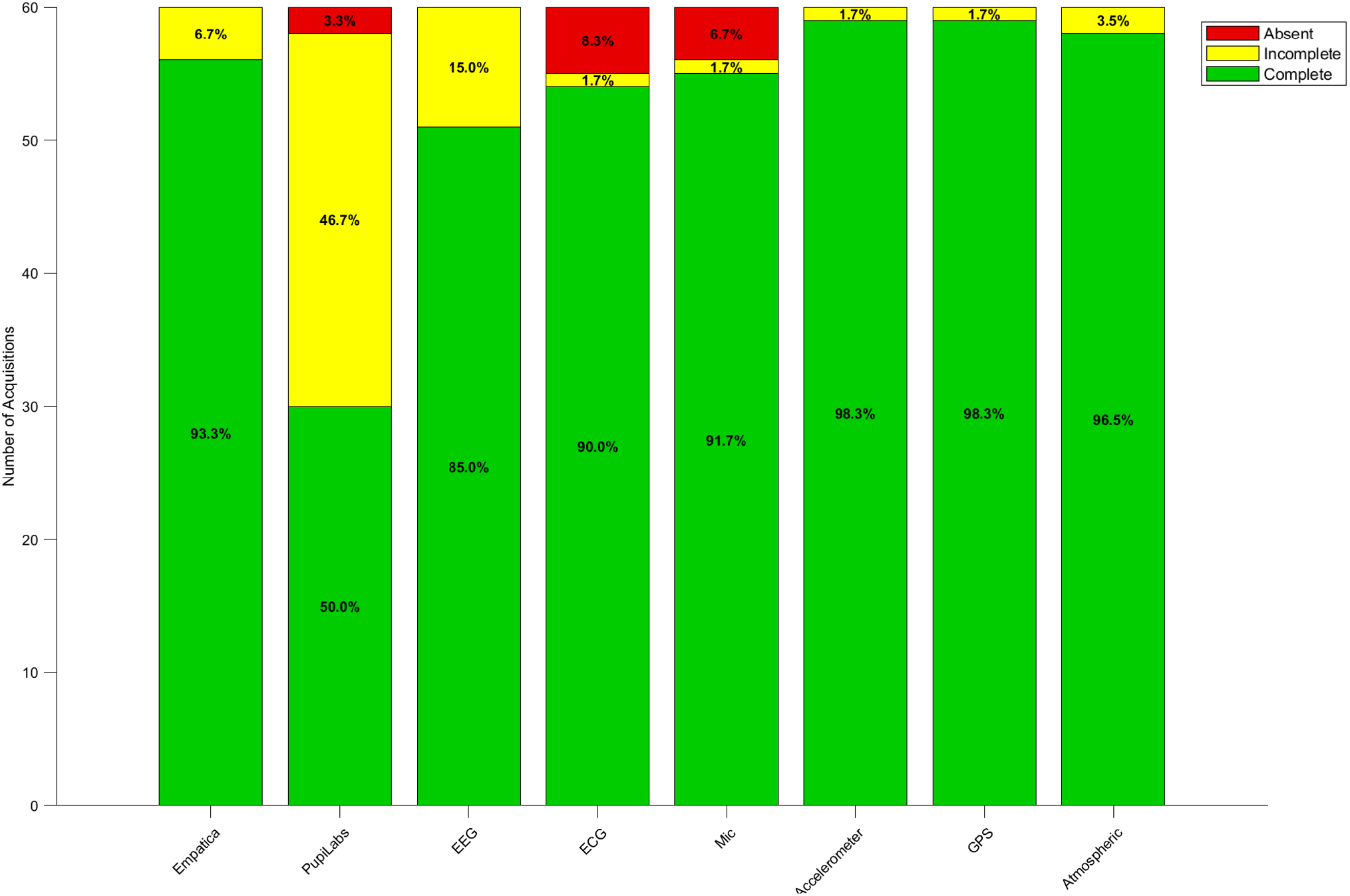
Sensor recording information across the 60 data acquisitions done in Lisbon. A specific sensor can be: complete, if during it recorded data during the whole path; incomplete, if it recorded data only until some part of the path; absent, if it recorded no data for that path.

### 3.7. Output data structure

Each acquisition had its output data stored in a folder containing all raw sensor data excluding EEG, which is saved independently in another directory by the NIC software, and added to the former posteriorly. The complete sensor data folder is then made available for analyses via a Python notebook, freely available in GitHub (https://github.com/emotional-cities/notebooks), that joins all the data in a single data structure. This structure is a spatiotemporal indexation of all the data streams. **Figure 5** illustrates how the different data streams, e.g. from climate sensors, derived from a single dataset can be plotted against the corresponding GPS coordinates. The set of GPS coordinates defining a walk is the common point between between-subject acquisitions and, as such, this framework allows the performance of analysis at the session-level and subject-level.

**Figure 5.**
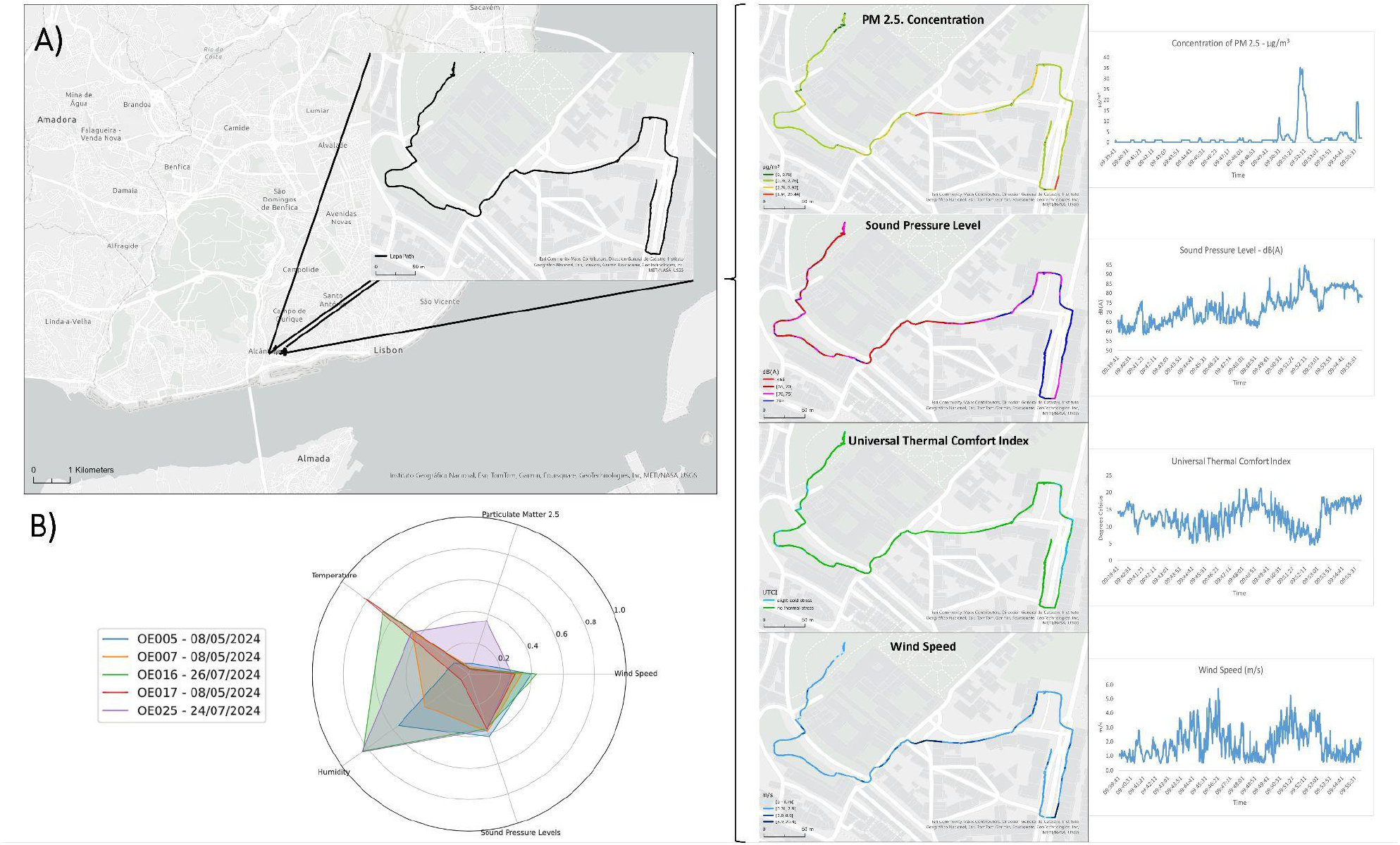
**A)** Different types of environmental data can be extracted from a single dataset corresponding to a specific path. On the right side is shown particulate matter with 2.5 μm, sound pressure level, Universal Thermal Comfort Index, and wind speed. Each of these metrics is plotted on top of the acquired GPS data. **B)** Radar chart showing normalized values of climate sensor metrics from the same path across different participants, showcasing the variability between sessions.

## 4. Discussion

One of the main advantages of having a powerful computer worn as a backpack is the possibility of acquiring data via wired connections, which mitigates data loss in wireless connections such as via Bluetooth. Furthermore, this novel wearable apparatus is highly modular and prioritizes open-source hardware and software which endow freedom to the researcher, a characteristic which often lacks in the outdoor EEG studies within Neurourbanism [37]. One technical aspect that stands out is the use of the HARP devices that allow the synchrony of devices sampling at different rates, which is a major challenge when merging different sensors together [38]. Notably, a pair of fully charged batteries lasts for 2 hours of continuous monitoring, besides the internal battery of the computer, allowing for a wide variety of tasks of variable duration. Finally, it is possible to use this backpack in virtual reality (VR) tasks and, if necessary, the climate station can be removed, giving more freedom of movement to the participant. The extensive port selection of the computer brings the added benefit of compatibility with external devices such as VR headsets and VR-accelerated computers capable of running demanding VR modelling tasks, thereby allowing recording while connected to another machine.

Due to its characteristics, EEG provides a bridge between mental health and urbanism, unlike any other neurophysiological tool. According to recent recommendations for mobile EEG outlined in [37], the Enobio’s 32-channel EEG system excels in providing a lightweight amplifier, a cap with standard electrode positioning, and freedom of analysis, including integration with the LSL protocol. The main disadvantages are mainly felt due to the demanding task, and include the presence of cable sway, the use of mastoid references, which are prone to go loose during a walk, and a lower channel count than the minimum recommended value of 64 channels. It should be noted that although dry electrodes were used, Enobio’s EEG system can also be used with gel electrodes. Nonetheless, the modularity aspect of the eMOTIONAL Cities Walker allows the quick and seamless integration of other, more suitable, EEG systems (see [39,40] for reviews).

Although the outdoor paradigm aims to simulate a natural walk, participants are inevitably affected by various sources of discomfort, including the backpack’s weight, the sensation of EEG electrodes, and the social exposure in public spaces where photos may be taken. Moreover, most sensors are sensitive to movement, leading to degraded signal quality. EEG, in particular, is highly susceptible to motion artefacts, with signal distortions that often exceed the amplitude of actual brain activity. Mitigation strategies include instructing participants to maintain a steady gait, using electrodes designed for better hair penetration, and using EEG systems with more channels that benefit more from data preprocessing tools.

Despite these challenges, the eMOTIONAL Cities Walker Backpack effectively integrates complex, multi-modal data streams essential for investigating brain–environment interactions in naturalistic settings. A key strength of the system lies in its use of redundancy to increase reliability and safeguard against individual sensor failure. Among all data types, spatiotemporal streams are the most critical, as they provide the framework for synchronizing all other signals. GPS plays a central role in spatial referencing, though its accuracy can be compromised in dense urban environments such as tunnels or areas with tall buildings. In such cases, video footage from the eye-tracking device’s external camera offers a valuable fallback for inferring spatial context. However, temporal synchrony—ensured by HARP devices—is irreplaceable, as it anchors all data streams to a precise timeline. Looking ahead, this apparatus sets a new benchmark for mobile neuroscience platforms, and future developments should prioritize enhanced comfort and discretion to further increase ecological validity and participant compliance in outdoor research.

## Notes

### Competing Interest Statement

The authors have declared no competing interest.

